# Pleiotropy constrains the evolution of immune system plasticity while promoting domain modularity

**DOI:** 10.64898/2026.05.27.728286

**Authors:** Danial Asgari, Ann T. Tate

**Affiliations:** Department of Biological Sciences, Vanderbilt University, Nashville, TN, USA; Evolutionary Studies Initiative, Vanderbilt University, Nashville, TN, USA

**Keywords:** Domain-specific evolution, Drosophila immunity, Gene expression evolution, evolutionary immunology, immune response, signaling protein evolution

## Abstract

Pleiotropic genes control multiple traits. This can result in evolutionary antagonism because adaptation that favors one trait can interfere with the function of another. While pleiotropic genes show statistical signatures of evolutionary constraint, many of them contain multiple domains that may evolve under different selective pressures. This could either strengthen or alleviate gene-level constraint. Here, we study pleiotropy within the immune system of six *Drosophila* species to disentangle gene and domain-level evolution. We hypothesized that the multifunctional nature of pleiotropic genes may promote within-gene variation in evolutionary rates of their domains compared to non-pleiotropic genes. Consistently, we found a greater within-gene variation in evolutionary rate among domains of pleiotropic genes than other gene classes, despite relatively low between-gene variation in evolutionary rates among pleiotropic genes. Non-pleiotropic genes, on the other hand, show a more heterogeneous selective pressure at the gene level. Regardless of pleiotropy status, domains within antiviral proteins show elevated evolutionary rates, while signaling protein domains show elevated ratios of radical to conservative amino acid substitutions, which likely have a significant effect on protein structure and function. Finally, an examination of plasticity in infection-induced gene expression responses across species revealed that non-pleiotropic genes with elevated evolutionary rates were also more likely to demonstrate variation in plasticity, but this relationship did not extend to pleiotropic genes. Overall, our results identify differences in evolutionary patterns across various biological levels (i.e., gene, domain, protein, and expression), showing that domain-specific evolution can potentially alleviate gene-level constraints.

## Introduction

Organisms have evolved immune defenses to combat microbial infections. Pathogens, in turn, constantly evolve new strategies to evade host defenses, giving rise to a continuous arms race between hosts and pathogens. This arms race is a major force behind the evolution of immune genes, which is driven either by adaptation through directional selection (Shultz and Sackton 2019; Nandakumar et al. 2025) or maintenance of polymorphism through negative frequency-dependent balancing selection (Unckless et al. 2016; Unckless and Lazzaro 2016; Minias and Vinkler 2022). Thus, it is surprising that many immune genes evolve under purifying selection (Jiggins and Kim 2006; Mukherjee et al. 2014). Previous studies have linked variation in evolutionary rates among immune genes to factors such as whether their products directly interact with pathogens (Sackton et al. 2007) and to participation of immune genes in other pathways (i.e., pleiotropy) (Williams et al. 2023). Disentangling these effects is crucial in understanding the evolution of immune responses.

Immune genes are disproportionately pleiotropic (Sivakumaran et al. 2011; Martin and Tate 2025), meaning their proteins have functions beyond immunity. For example, the cytokine IL-6 in humans and its homolog in *Drosophila melanogaster*, unpaired 3 (Upd3), have both immune and developmental functions (Hombría et al. 2005; Kishimoto 2006; Jenkins et al. 2021). Perhaps the best-known pleiotropic signaling pathway in *D. melanogaster* is Toll signaling, which is involved in both the production of small antimicrobial peptides (AMPs) and in dorso-ventral patterning during early development (Valanne et al. 2011). How does pleiotropy within immune systems affect the evolutionary arms race between hosts and pathogens?

Fisher’s geometric model suggests that for complex traits, mutations with large effects are more likely to be deleterious, thus implying evolutionary constraint on pleiotropic genes (Fisher 1930). Pleiotropic constraints have been observed across diverse species and contexts (Mank et al. 2008; Papakostas et al. 2014). Therefore, it is reasonable to assume that pleiotropy hampers the adaptation of immune genes to novel pathogens. Williams et al. (2023) explicitly tested this hypothesis across the *Drosophila* clade and showed that immune genes that are pleiotropic with development evolve more slowly than non-pleiotropic immune genes. They also explored how pleiotropy varies across functional classes, including signaling proteins, receptors, and effectors. Proteins involved in immune signal transduction (e.g., in Imd and Toll signaling) tend to be more pleiotropic and, thus, evolve more slowly.

In addition to imposing a stronger purifying selection on coding sequences, pleiotropy may also constrain gene expression, although the evidence for this effect is mixed. One study examining adaptation to heat stress in *Tribolium* beetles found that gene expression changes during evolution largely occur through indirect selection acting on highly pleiotropic eQTLs, which affect the expression of many other genes (Koch et al. 2025). In contrast, another study found that highly pleiotropic genes, identified by the number of protein-protein interactions, did not change in expression upon thermal adaptation of freshwater salmonid fish populations (Papakostas et al. 2014). The authors suggested that expression of pleiotropic genes is constrained because changes that benefit one function may be harmful to others. These conflicting results might come from differences in the systems studied or the definition of pleiotropy. Hodgins et al (2016) showed that, in conifers, genes with greater variability in expression plasticity in response to abiotic stress evolve more rapidly. This was possibly driven by relaxed selection on genes with less conserved expression patterns. However, they did not consider the effect of pleiotropy on responses to biotic stressors (e.g., infection). Immune genes often show plastic expression responses to infection. Therefore, to get a clearer picture of immune system evolution, it is crucial to examine how evolutionary rates vary among genes with different levels of induction across species, and how pleiotropy shapes both gene induction and sequence evolution. Addressing this missing piece of the puzzle can provide insight into why evolutionary rates vary so widely among genes involved in immune responses.

Most proteins have multiple domains (Wang and Caetano-Anollés 2006), and pleiotropic proteins interact with other entities (e.g., other proteins or DNA) through these domains. Thus, a stronger evolutionary constraint is expected on domains compared to the entire protein.

However, most studies of domain evolution focus on domain architecture (i.e., the order of domains in the protein) (Vogel et al. 2004; Fong et al. 2007; Basu et al. 2008; Buljan and Bateman 2009; Forslund and Sonnhammer 2012), or domain-specific evolution rather than comparing the evolution of the domain to the full-length genes. One study did suggest that pleiotropic interactions can affect evolutionary constraints on domains. Specifically, transcription factor-DNA interactions constrain DNA-binding domains (slower evolution), whereas transcription factor-transcription factor interactions constrain the evolution of the entire protein (Chesmore et al. 2016). Another study by Mannakee and Gutenkunst (2016) found that domains with more dynamical influence on signaling evolve more slowly, and this effect is stronger for proteins with a higher degree of connectivity (i.e., pleiotropic proteins). A genome-wide study in *D. simulans* found higher rates of adaptive substitution in non-domain regions of proteins, but did not consider immune genes specifically (Holloway and Begun 2007). Domain-specific studies of immune gene evolution are scarce or focus on a single gene. For example, Begun and Whitley (2000) found strong adaptive substitution in the regulatory region of Relish in *D. simulans*, a protein responsible for the induction of AMPs in Imd signaling, which is possibly driven by host-pathogen co-evolution. Currently, we do not know whether such patterns exist across proteins that perform different functions in signaling pathways.

Given that immune genes are highly pleiotropic with traits under strong purifying selection, such as development and the nervous system (Senthilkumar et al. 2026), the immune system provides an excellent opportunity to address this gap. We hypothesize that domains within pleiotropic genes evolve under a heterogeneous selective pressure due to the multifunctional nature of these genes. In contrast, we expect that in less constrained genes, such as non-pleiotropic immune genes or those that directly interact with pathogens (e.g., involved in pathogen detection or killing), genes and their domains would evolve at similar rates. We predict a similar pattern at the protein level, where replacement of amino acids by those with dissimilar biochemical properties is more likely for less functionally constrained genes and domains. Finally, we expand our analysis beyond sequence-level evolution and examine whether pleiotropic constraint also extends to gene expression regulation. Our results have implications beyond the immune system, as they provide general insights into how pleiotropy and functional constraint shape domain and gene evolutionary dynamics.

## Results

### Pleiotropy constrains evolution at both the gene and domain levels while promoting within-gene variation in domain evolutionary rates

We tested for pleiotropic constraints by estimating dN/dS for the four classes of genes: randomly selected genes (hereafter ‘random genes’), pleiotropic immune genes (hereafter ‘pleiotropic’), non-pleiotropic immune genes (hereafter ‘immune genes’), and developmental genes. Pleiotropic genes and their domains evolve more slowly than immune genes and more closely resemble developmental genes (Fig.S3). Immune genes have a similar rate of evolution to randomly selected genes at the gene level; however, they evolve faster at the domain level (Fig.S3).

To examine the relationship between gene and domain evolution across the four groups, we measured the relationship between domain-level dN/dS and gene-level dN/dS (Fig. 1A) by using a linear mixed-effects model (Equation 1). We observed that in randomly selected genes, domain dN/dS closely matches gene dN/dS (β = 1.0). In other gene classes, domain dN/dS increases more slowly than gene dN/dS (Table S1). We also fit Equation 3 simultaneously on all four classes to examine whether this difference is significant. We found that the pleiotropic and developmental classes have significantly smaller slopes than the immune and random genes (Table S2).

**Fig 1.**
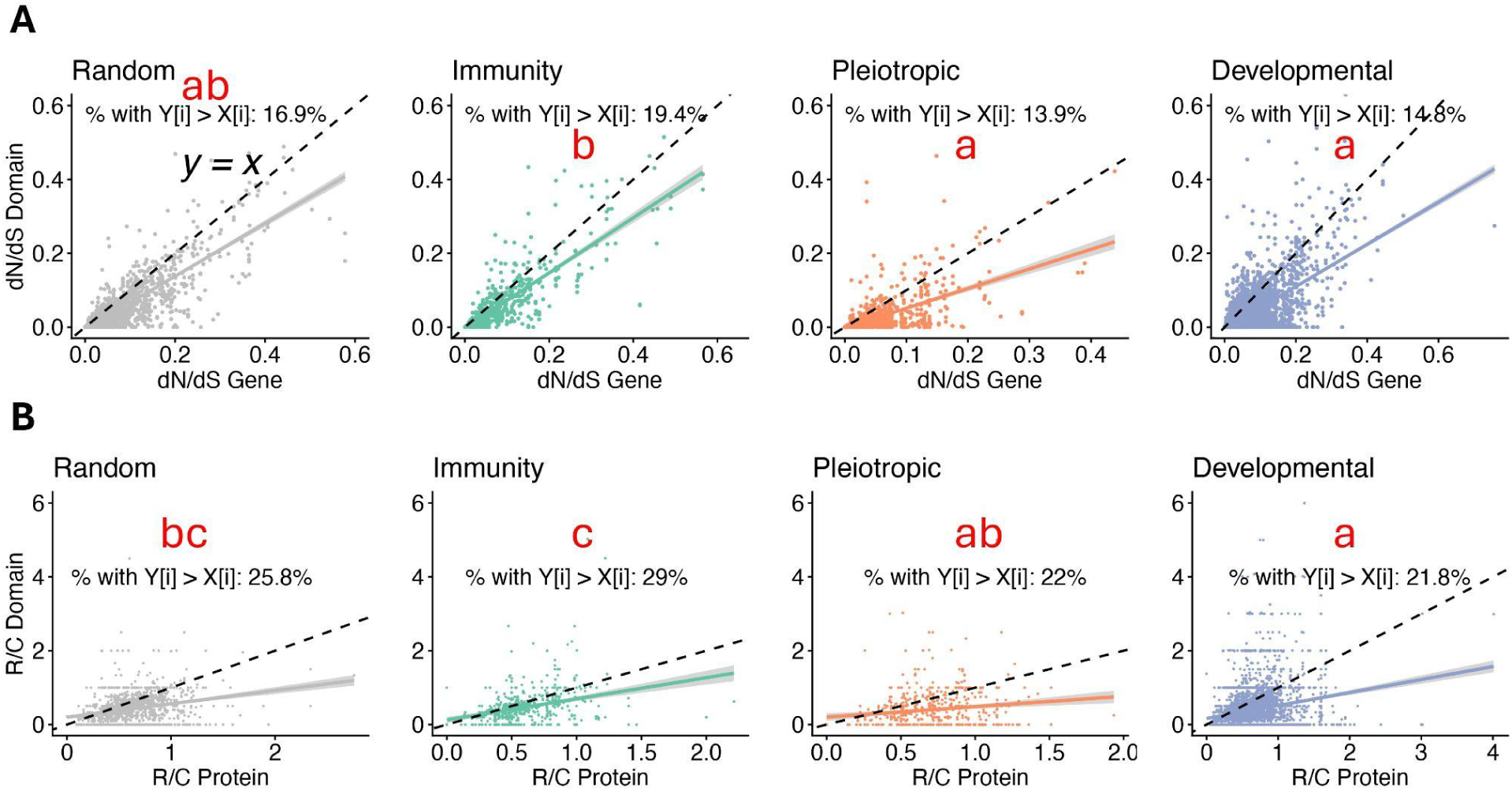
Domains and genes show more similar dN/dS and R/C in immune genes (regression line close to y = x) compared to pleiotropic and developmental genes. Panel A shows dN/dS for genes (X-axis) and domains (Y-axis) across three gene classes. The percentage of genes with higher dN/dS in domains than in genes is shown in each plot. Panel B shows the proportion of radical/conservative (R/C) substitution across the species tree for proteins (X-axis) and domains (Y-axis). Groups sharing a letter have similar proportions of Y[i] > X[i] based on pairwise tests for equality of proportions.

To quantify similarity between and within genes (i.e., between domains), we used the intra-class correlation coefficient (ICC) (Equation 2). A small ICC indicates either decreased between-gene variance and/or increased within-gene variance. This parameter was largest for random genes (ICC ≈ 0.80), moderate for immune (ICC ≈ 0.40) and developmental genes (ICC ≈ 0.38), and smallest in pleiotropic genes (ICC ≈ 0.20). These differences were significant, as ICC values were lower in the pleiotropic group in ∼100% of bootstrap samples (Fig.S4). Pleiotropic genes have the lowest between-gene variance, while their within-gene variance was elevated and close to that of immunity genes (Fig.S5A). This suggests that pleiotropic genes show high heterogeneity in evolutionary rates among domains despite low divergence between genes.

Williams et al. (2023) showed that among recognition, signaling, and effector immune classes, signaling contains the highest proportion of pleiotropic genes, and we therefore predicted that the domains from pleiotropic genes would be enriched for signaling functions relative to other gene classes. We tested this hypothesis by performing enrichment analyses for pleiotropic, immune, and developmental domains relative to domains of randomly selected genes. Consistent with our hypothesis, pleiotropic genes were significantly enriched for domains involved in signaling (Table S3), including protein kinase and small GTPase domains (e.g., Ras and ADP-ribosylation factor families). Developmental genes showed enrichment for a functionally diverse set of domains, such as those involved in transcription, structural functions, protein-protein interactions, and signaling. As expected, domains in immune genes were enriched for host defense activity. We next asked whether high within-gene variance in domain evolutionary rates is specific to pleiotropic genes or simply driven by domains commonly found in them, such as kinases. To this end, we identified genes with at least one kinase domain across the four gene classes. This included 40 pleiotropic genes, 9 immune genes, 155 developmental genes, and 25 random genes. We then used Equation 1 to examine the relationship between domain and gene-level dN/dS and calculated ICC for each class. We found that ICC was ∼10 times lower in pleiotropic genes compared to immune and developmental genes (Table S4). Pleiotropic genes with a kinase domain had the lowest between-gene variance, and within-gene variance was still elevated, similar to developmental genes (Fig.S5B). Thus, pleiotropic genes, unlike immune genes, maintain the same pattern, namely the smallest between-gene variance and elevated within-gene variance, even after controlling for domain type (e.g., kinases).

To predict how non-synonymous substitutions affect protein structure, we counted radical and conservative amino acid substitutions across the phylogenetic tree for genes and their domains and calculated the ratio of radical to conservative substitutions (R/C). Radical substitutions are less likely to occur, as they entail the replacement of an amino acid with one that has different physicochemical properties; thus, a higher R/C ratio likely reflects changes in protein structure. The relationship between gene-level and domain-level R/C was weaker than the one observed for dN/dS: immune genes, β = 0.57, pleiotropic, β = 0.30; and random and developmental β = 0.37. In addition, the range of ICC was lower than ICC for dN/dS estimates (ICC ≈ 0.07–0.13, with the highest value in immune genes), which is driven by very small within-gene variance across gene classes (Fig.S6). The pleiotropic group had the second-highest within-gene variance, with developmental genes showing the highest variance.

We also calculated the proportion of domains that evolve faster than their corresponding genes (data points above y = x in Fig.1A). We found that this proportion is significantly higher for immune genes (∼20%) than pleiotropic and developmental genes (∼14%). The randomly selected genes were not different from the other categories (Fig.1A). Next, we examined whether differences in dN/dS among domains and genes held for R/C as well, and found a similar pattern between R/C and dN/dS (Fig.1B). Specifically, immune genes show a significantly larger proportion (∼29%) of domains with a higher R/C than the full-length protein compared to all other groups. This ratio was lower in pleiotropic and developmental genes (∼22%). For randomly selected genes, the ratio (∼26%) was close to that of immunity and pleiotropic classes but higher than that of developmental genes.

To further examine the strength of selection, we estimated the rate of non-adaptive substitution (ω_na_), which reflects the tolerance of evolution for maintaining weakly deleterious mutations, across the four classes of genes in three populations of *D. melanogaster* (Raleigh, Munich, and Barcelona) from the *Drosophila* Evolution over Space and Time (DEST) dataset (Kapun et al. 2021; Footprints of worldwide adaptation in structured populations of *Drosophila melanogaster* through the expanded DEST 2.0 genomic resource 2025). We found that across all three populations, immune genes have a higher ω_na_ than pleiotropic genes, while developmental genes show the lowest rate. In Munich and Barcelona populations, ω_na_ is higher in random genes than in immune genes, whereas the opposite pattern is observed in Raleigh. The results for the Raleigh population are shown in Fig.2 (for all three populations, see Fig.S7). At the domain level, pleiotropic genes have the lowest ω_na_ across all populations.

**Fig 2.**
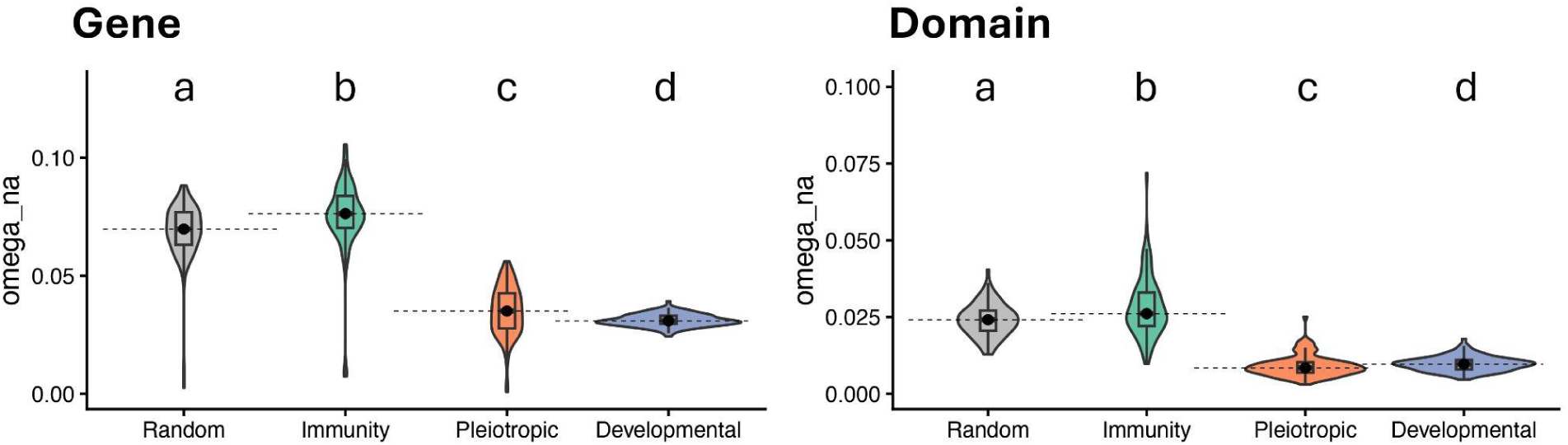
Pleiotropic immune genes have the second-lowest rate of non-adaptive substitutions at the gene level and the lowest at the domain level. The Y-axis shows the rate of non-adaptive substitution (ω_na_) across three classes of genes (X-axis) for the Raleigh population. The left panel shows the results at the gene level and the right panel at the domain level. All comparisons are significant (Wilcoxon test). To improve visualization, medians are extended with black dashed lines to facilitate comparison.

Developmental genes show slightly but significantly higher ω_na,_ and immune or randomly selected genes have the highest ω_na,_ depending on the population (Fig.2). We also estimated the rate of adaptive substitution (ω_a_) and the proportion of adaptive substitution (⍺) at the gene level (Fig.S8). The rate of adaptive substitution (ω_a_) did not show a consistent pattern across the populations, with similar values across all four gene classes. The proportion of adaptive substitutions (⍺) was highest in developmental genes, followed by pleiotropic genes. Depending on the population, either random genes or immune genes had the lowest ⍺.

### Regardless of pleiotropic status, domains within immune proteins show distinct patterns of divergence and amino acid change across different functional classes

The function of a gene within the immune system, regardless of its pleiotropic status, can shape its evolution. Previous studies have shown that within immune pathways, evolutionary rates vary across different functional classes, such as pathogen recognition, signaling, and effector (killing) (Han et al. 2013). We examined this by estimating dN/dS and R/C ratios for genes and their corresponding domains in two major signaling pathways of *D. melanogaster*, namely Imd and Toll signaling. We found that most genes in both pathways have domains that evolve more slowly than their corresponding gene. A few genes, however, have domains that evolve faster than the full-length gene (labeled in the plots in Figure 3). For example, the ankyrin domain of Relish in Imd, and the dimerization and DNA-binding domains of Dif in Toll, evolve faster than their respective genes. The R/C ratio provided additional insight into the evolution of protein structure. For example, although the dimerization and DNA-binding domains of Relish evolve slowly (smaller dN/dS than the gene), the proportion of radical to conservative changes in these domains is higher than in the full-length protein. We also observed high R/C ratios in the domains of some pathogen recognition proteins in both Imd (PGRP-SD) and Toll (GNBP3) signaling pathways.

**Fig 3.**
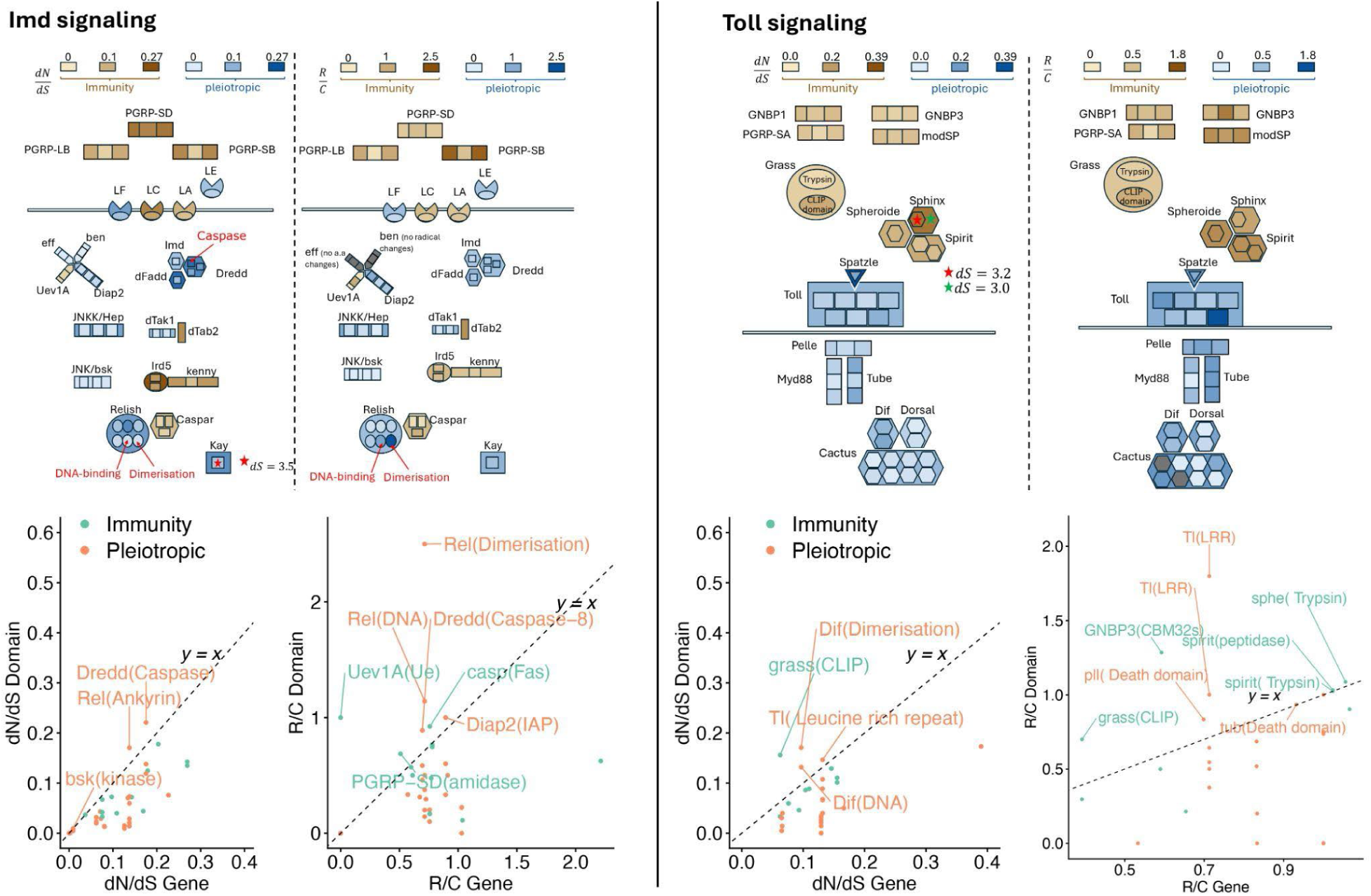
dN/dS and R/C for genes and their domains for components of Imd (left) and Toll (right) signaling. Shades of blue show the magnitude of dN/dS and R/C for pleiotropic genes and domains. Shades of brown show the magnitude of dN/dS and R/C for non-pleiotropic immune genes and domains. Small plots on the bottom show the dN (or R) on the Y-axis over dS (or C). The genes (or proteins) with larger dN/dS (or R/C) for domains than the entire gene (or domain) are labeled on the plots. The genes and domains with slightly inflated dS (dS>3) are marked by a star. This figure is adapted from Williams et al. (2023).

As expected, genes whose products interact with pathogens (recognition, effector, and antiviral genes) have a higher median dN/dS than signaling and “other” genes (Fig.4A, Table S5). Antiviral genes were not significantly different from other categories at the gene level but did have significantly higher dN/dS than all other categories at the domain level (Fig.4B).

**Fig 4.**
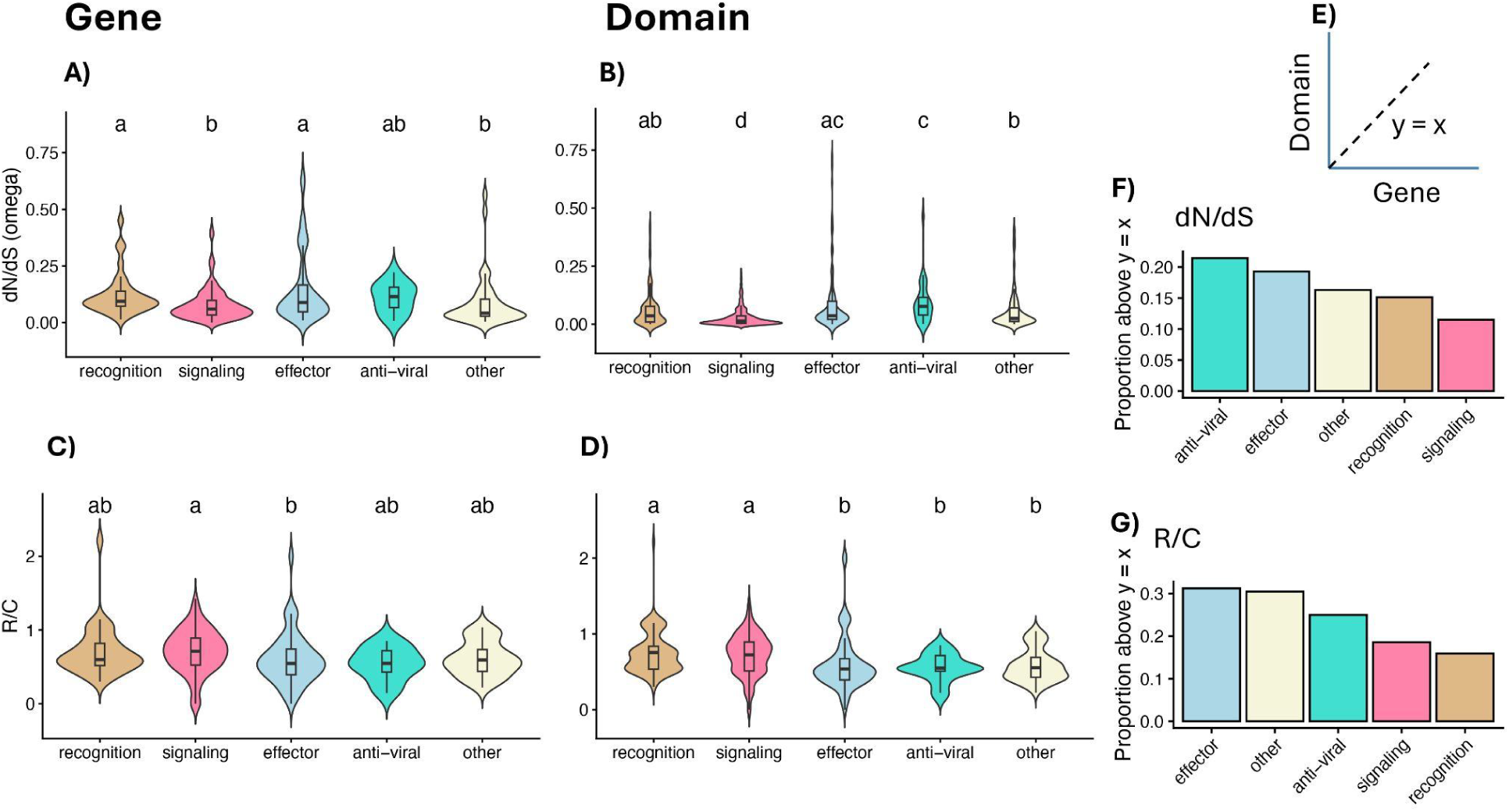
Rapid change in immune genes that directly interact with pathogens. Comparison of different classes of immune classes (X-axis) using dN/dS across genes (A) and domains (B), and R/C ratios across genes (C) and domains (D) is shown. The right-most column shows the proportion of genes with higher dN/dS (F) and R/C (G) in the domain regions compared to the rest of the gene (or protein). Groups sharing a letter are not significantly different (Wilcoxon test).

Effector genes also have higher dN/dS at the domain level compared to signaling and “other” genes. We conducted the same analysis for R/C. At the protein level, signaling proteins have significantly higher R/C than effectors (Fig.4C and Table S5), and at the domain level, recognition and signaling classes have significantly higher R/C than other classes (Fig.4D and Table S5). We also plotted dN/dS (and R/C) for genes (and proteins) on the X-axis and for domains on the Y-axis to determine the proportion of genes with higher dN/dS or R/C at the domain level compared to their corresponding genes (Fig.4E). The Antiviral class has the highest proportion of genes with domains evolving faster than the corresponding genes (Fig.4F). The effector class has the highest proportion of genes with higher R/C at the domain level than the corresponding protein (Fig.4G). For recognition proteins, the proportion of fast-evolving domains, compared to the full-length genes, or with higher R/C than the corresponding protein, is generally low (second lowest for dN/dS and lowest for R/C).

### Pleiotropy constrains variation in plastic response to infection, whereas in non-pleiotropic immune genes, sequence divergence is correlated with variation in plasticity

Some previous studies have shown that pleiotropy constrains only sequence-level evolution (Molodtsova et al. 2014), while others suggest it affects both sequence and gene expression (Papakostas et al. 2014; Koch et al. 2025). To investigate how pleiotropy influences the immune response, we took advantage of RNA-seq data from Salazar-Jaramillo et al. (2017) to examine the relationship between dN/dS and induction of immune genes following exposure to parasitoid wasps for four out of the six species within the *Drosophila* clade used in our analysis. We measured changes in gene expression as log_2_ fold changes (LFCs) for orthologous genes and constructed PCAs for pleiotropic, non-pleiotropic, and random genes (with the random sample size equal to the average of the other two groups) based on LFCs following exposure to the parasite. We observed low dispersion for pleiotropic genes compared to non-pleiotropic and random genes (Fig.5A), indicating smaller inter-species variation in transcriptional response for pleiotropic immune genes. Low dispersion on PCA suggests similar LFCs for pleiotropic genes. To investigate the magnitude of LFCs, we plotted the average LFC across orthologous genes for the three gene classes. We found that pleiotropic immune genes have the highest density at zero, while non-pleiotropic immune genes show the heaviest tails (more genes with large changes in expression) (Figure 5B). The LFC distribution for random genes fell between the two other distributions (Figure 5B). To quantify the association between divergence at the sequence and expression level, we then measured the correlation between the distance from the PCA centroid and dN/dS (dS < 3). We found a weak but significant positive correlation between dispersion on PCA and dN/dS across all three gene classes (Fig.S9). The correlation was strongest for random genes, moderate for non-pleiotropic immune genes, and weakest for pleiotropic genes. Next, we classified genes as “central” or “scattered” (Figure 5C) based on their distance from the centroid (using thresholds from the closest 5% to 50% of genes). We then compared the median dN/dS between central and scattered genes for both pleiotropic and immune genes. For random genes across most threshold values, and for non-pleiotropic immune genes at larger threshold values, the dN/dS of scattered genes becomes significantly larger than that of central genes; however, this was not true for pleiotropic genes (Fig.5D). We next asked whether the larger dN/dS values for scattered non-pleiotropic immune genes are driven by relaxed selection or by positive selection. We found that in the Munich population, ⍵_a_ and ⍺ values were significantly larger for scattered genes at 50% threshold compared to central genes (Fig.S10). ⍵_na_ was also slightly larger for scattered genes. Overall, this analysis suggests that pleiotropy constrains the magnitude and variation in plastic immune response to infection across species of *Drosophila.* In non-pleiotropic immune genes, elevated dN/dS, associated with positive selection, is correlated with a greater variation in gene expression response.

**Fig 5.**
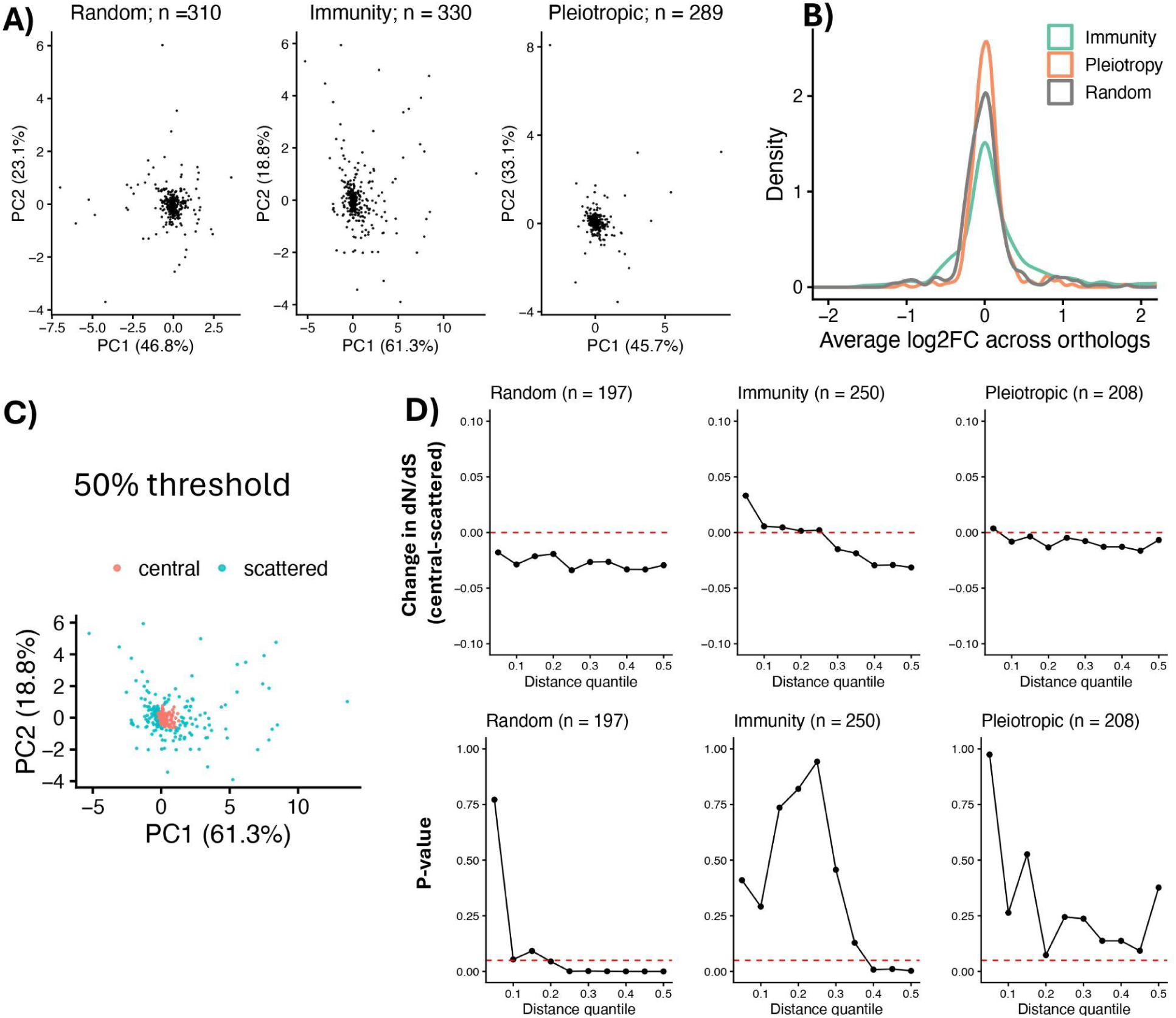
Greater plasticity is associated with elevated dN/dS in non-pleiotropic immune genes. Panel A shows PCA plots for log_2_ fold change values across orthologous genes of four *Drosophila* species for random genes (left), immune genes (center), and pleiotropic genes (right). The number of genes for which LFCs are available is reported above each plot. The distributions of LFCs across three classes are shown in panel B. The X-axis shows the magnitude of LFCs, and the Y-axis shows the density. An example of the classification of genes into central and scattered groups is shown in panel C. The top row in panel D compares the median dN/dS between central and scattered genes across thresholds ranging from the closest 5% to 50% of genes. The dN/dS values here are only reported for dS < 3; hence, the difference in the number of genes between panel A and D. In the top panel, the red dashed line indicates zero difference. The bottom row shows the p-value for the difference in dN/dS median. In the bottom panel, the red dashed line indicates the significance threshold (p = 0.05).

## Discussion

By examining the immune system of a *Drosophila* clade, we showed that domains of pleiotropic genes, similar to developmental genes, are under strong purifying selection. Pleiotropy not only constrains sequence evolution but also, at the gene expression level, constrains the plastic response to infection. Despite these constraints, pleiotropic genes show greater variation in evolutionary rates across their domains compared to other gene classes (Fig.S5), indicating modular evolution at the domain level. On the other hand, non-pleiotropic immune genes generally have similar gene and domain-level evolutionary rates and are more likely to contain domains that evolve faster than their genes (Fig.1). We observed a greater sequence divergence in non-pleiotropic immune genes that show a stronger variation in plastic response to infection across species. This suggests that in the absence of pleiotropic constraint, immune genes diverge both at the coding and regulatory levels. Regardless of the pleiotropic status of a gene, we found rapid evolution of domains within genes that interact with pathogens, such as effectors and anti-viral genes. By examining individual genes within immune signaling pathways, we found that domain evolution at both the nucleotide and protein levels is context-dependent and shaped by the domain function.

Pleiotropy constrains the evolution of a gene, and this should be more readily observed at the domain level because domains carry out the fundamental functions of a gene. Consistent with this prediction, our analyses show that at the domain level, pleiotropic immune genes evolve more slowly than non-pleiotropic ones and have a similar rate of evolution to developmental genes (Fig.S3). Notably, the domains of pleiotropic genes have an even lower tolerance than developmental genes for accumulation of weakly deleterious mutations (Fig.2 and Fig.S7). Even with this overall constraint, pleiotropic genes must perform at least two different functions (e.g., immunity and development); thus, it is reasonable to assume that this constraint could be alleviated by being distributed among domains. Consistently, we found more variation in dN/dS estimates between domains of pleiotropic genes compared to non-pleiotropic immune genes and developmental genes. Pleiotropic domains were enriched for cell signaling functions, but examination of kinases across gene classes showed elevated within-gene variation in pleiotropic genes, suggesting that differences among gene classes are not simply confounded by the types of domains within them. It is possible that the modular nature of proteins involved in signaling, in which different domains perform distinct biochemical functions (Lee and Yaffe 2016), such as adding (e.g., phosphorylation), removing (e.g., phosphatase activity), or recognizing post-translational modifications (e.g., protein-protein interaction domains), may underlie the variation in evolutionary rates observed across domains within pleiotropic genes.

There could be other explanations for why domains evolve at different rates. Within multidomain proteins, younger domains evolve faster than older ones (Toll-Riera and Albà 2013). Thus, combining domains with different ages may result in higher variance in evolutionary rates of domains within a gene. Martin and Tate (2025) found that pleiotropic genes tend to be middle-aged or ancient rather than young. Thus, if domains and genes have similar ages in pleiotropic genes, this hypothesis is unlikely. Another more likely explanation comes from a study by Zhou et al. (2008). They showed that within the same protein, domains with higher contact density (average contacts per residue) evolve faster. It is possible that within signaling pathways, pleiotropic proteins use domains with different contact densities to efficiently transmit information, thus resulting in variation in evolutionary rates. Heterogeneous selective pressure on the domain of pleiotropic genes might be caused by the division of tasks among domains. Alternatively, domains that participate in crucial functions (e.g., protein-protein binding) could be under stronger purifying selection, while domains that perform peripheral functions evolve under weaker constraints. Unfortunately, there is still not enough fine-grained genome-wide information about domain-specific functions to achieve adequate power to test these hypotheses.

Although the domain-level evolutionary rate in pleiotropic immune genes is generally constrained, some domains evolve faster than their genes. In Imd signaling, for example, we found that the caspase domain of Dredd and the ankyrin domain of Relish evolve faster than corresponding genes (Fig.3). Dredd mediates the caspase-mediated cleavage of Relish (Stoven et al. 2003), following which Relish is split into two fragments: ankyrin (the inhibitory domain), which remains in the cytoplasm, and the Rel homology domain, which enters the nucleus to transcribe AMPs (Stöven et al. 2000). Sackton et al. (2007) showed that positively selected sites in Dredd are located within the caspase domain. Begun and Whitley (2000) found adaptive substitutions in the ankyrin domain of *D. simulans* and suggested that pathogens that interfere with cytoplasmic signaling via the type III secretion system might promote adaptive evolution for such signaling proteins. For example, *Yersinia pestis,* the causative agent of plague, uses the type III secretion system to disrupt NF-κB signaling in mammals (Viboud and Bliska 2005). This host-pathogen co-evolution hypothesis might also explain the elevated evolutionary rate for some Toll signaling transcription factors, such as the dimerization and DNA-binding domains of Dif (Fig.3). While previous studies have provided ample evidence of arms races in recognition and effector proteins, our results suggest that domains within signaling proteins are not immune to arms races and still need to adapt despite the increased pleiotropic constraints on them.

We found that a subset of domains that evolved more slowly than the full-length gene at the nucleotide level showed a higher ratio of radical to conservative amino acid substitutions at the domain level compared to the full-length protein. This means that amino acid substitutions are more likely to be of the kind that change the structure and function of the protein at the domain level compared to the full-length protein. Thus, although the evolution of the domain is slow, the few substitutions that do occur tend to be more radical than conservative. For example, domains of some recognition proteins, such as PGRP-SD in Imd or GNBP3 in Toll, exhibit a higher proportion of radical to conservative substitution compared to the overall protein. This is consistent with a high ratio of R/C for recognition proteins (Fig.4D). This overall trend, and in particular for proteins like PGRP-SD and GNBP3, likely reflects host-pathogen co-evolution at the interface of host-pathogen contact, consistent with previous literature (Dhakad and Obbard 2026). Even though signaling proteins generally have lower dN/dS estimates, they show elevated R/C ratios, especially at the domain level. This is consistent with some previous studies that identified signatures of positive selection on signaling proteins in Imd and Toll pathways (Han et al. 2013). However, we must be careful in interpreting R/C, because it does not account for mutation biases or normalize by the number of radical and conservative sites, and therefore reflects observed substitution patterns rather than the evolutionary rate (Suzuki 2007).

We show that pleiotropic constraint not only affects coding sequence evolution but also plastic gene expression in response to infection (Fig.5). Several studies in plants, mammals, and birds have shown that gene expression divergence and dN/dS are correlated (Warnefors and Kaessmann 2013; Hodgins et al. 2016), indicating that selection acts on both sequence and expression. We observed this both for random genes and non-pleiotropic immune genes, but not for pleiotropic genes. Across *Drosophila* species, pleiotropic genes showed a smaller overall change and similar LFC following exposure to parasitoid wasps. Further, we showed that the elevated level of dN/dS in non-pleiotropic immune genes with greater variation in plasticity is associated with positive selection. These findings suggest that pleiotropy constrains plastic expression of immune genes across species in addition to imposing stronger purifying selection on their coding sequences. Zhong et al (2021) found a positive correlation between coding sequence divergence and basal gene expression divergence for pattern recognition receptors across two rodent species. However, in this study, pleiotropy mainly constrained sequence evolution and not gene expression. Importantly, they looked at basal gene expression and not induction following infection. Similarly, a study on social behavioral traits of honeybees observed decoupling of pleiotropic constraint, showing coding but not regulatory regions are affected (Molodtsova et al. 2014). Interconnected genes were reported to experience strong purifying selection at the coding level, while connectivity had no influence on the strength of selection acting on regulatory regions. By examining immune gene induction, we show that, unlike previous studies, pleiotropic constraint acts on both coding sequence evolution and plastic transcriptional responses following immune stimulation. Unfortunately, in the *Drosophila* clade, there are limited RNA-seq studies of bacterial or viral infections in closely related species that allow us to test for such connections. To our knowledge, this is the first study that identifies a link between pleiotropic constraints on both coding and induction of immune genes.

Overall, our study provides generalizable insight regarding the evolution of pleiotropic genes that can be extended to systems beyond immunity and potentially to other species.

Importantly, we showed that pleiotropy constrains evolution at both the domain sequence and expression levels, while simultaneously promoting variation in evolutionary rates across domains. Our results also provide a strong framework for understanding how pleiotropy and functional constraints shape domain evolution in pathways in which proteins perform diverse functions.

## Materials and Methods

### Identification of orthologous sequences

The immune system is highly pleiotropic with development. Specifically, about 40-44% of immune genes are pleiotropic with development (Williams et al. 2023). In addition, genes involved in early development are under strong purifying selection due to a higher number of interactions with other genes (Artieri et al. 2009). Thus, developmental pleiotropy within the immune system provides an excellent framework for studying evolutionary constraints. To begin our analyses of selection at gene versus domain levels, we took advantage of a previous dataset (Williams et al. 2023) that categorized a subset of insect genes into three classes: pleiotropic immune genes (n = 299), non-pleiotropic immune genes (n = 454), and developmental genes (n = 3,047). To provide a reference in the broader genomic context, we also added a set of randomly selected genes across the genome (Fig.S1). The number of randomly selected genes is equal to the average size of the three previous groups (n = 1,267). To categorize immune genes based on function and regardless of pleiotropy status, we used previous literature (Sackton et al. 2007; Early et al. 2017) and supplemental Table 2 of Williams et al. (2023), and divided pleiotropic and non-pleiotropic immune genes into recognition, signaling, effectors, antiviral, and “other” immune genes.

We used the NCBI database to identify orthologous genes in six species of Drosophila (*D. melanogaster, D. ananassae, D. erecta, D. sechellia, D. simulans,* and *D. yakuba*). To this end, we first used Ensembl Metazoa (release 62) to obtain NCBI gene IDs corresponding to FlyBase IDs. Then, we found the ID of the longest protein sequence for each NCBI gene ID. We downloaded these *D. melanogaster* protein sequences for all classes (random, pleiotropic, non-pleiotropic, and developmental), and blasted (BLASTP) them to the proteome of the five other species of *Drosophila*. We identified an orthologous protein as the best hit (highest bitscore), and filtered out low-quality blast results (bitscore < 100).

### Sequence alignment and dN/dS estimation for genes and domains

We downloaded the CDS of every orthologous gene (six or fewer CDS sequences per gene, depending on the number of identified orthologs) from NCBI, and for every gene, we translated CDS sequences to the protein sequence using SeqKit v2.10.1 (Shen et al. 2016). Protein sequence alignment was performed using MAFFT v7.525 (Katoh and Standley 2013). Next, we generated codon alignments of CDS sequences using PAL2NAL v14 (Suyama et al. 2006) based on the translated protein sequences. Codon alignments were trimmed using Gblocks v0.91b (Castresana 2000). Following Williams et al. (2023), species trees were pruned to retain only species with extant sequences in the codon alignments using nw_prune from the Newick Utilities package (Junier and Zdobnov 2010). To calculate the rate of non-synonymous to synonymous substitutions (dN/dS) across orthologous genes, we ran PAML v4.10.7 (Yang 2007) using the trimmed sequences and pruned species trees, considering only alignments with more than one species (model = 0, NSsites = 0, seqtype = 1, CodonFreq = 2, cleandata = 1).

We extracted both dN and dS values to assess synonymous substitution saturation (dS > 3) (Wolf et al. 2009; Hermansen et al. 2017; Northover et al. 2020), and filtered out saturated dN/dS estimates.

To identify the coordinates of functional domains, we used hmmscan from HMMER v3.4 to scan translated CDS sequences against the Pfam database (Paysan-Lafosse et al. 2025).

For better alignment, short domains (length < 20) were filtered out. We also removed domains with low confidence (E-value > 1 × 10⁻⁵). The coordinates of the domains were used to extract the corresponding protein and CDS sequences. These were then used as input for PAL2NAL for alignment and subsequently for PAML to estimate dN/dS, following the same approach as for the full-length gene described above. We measured dN/dS once using the species tree and another time using gene trees of orthologous genes, and found consistent results across the two measurements (Fig.S2). Thus, we used the species tree for all analyses presented here.

We ran linear mixed-effects models (Equation 1) to examine the relationship between domain-level dN/dS and gene-level dN/dS. In these models, protein identity is used as a random effect to account for similarities among domains from the same protein. We additionally calculated the intra-class correlation coefficient (ICC) to quantify how much variation is explained by differences among genes (Equation 2). These were also done for R/C measures (see below).

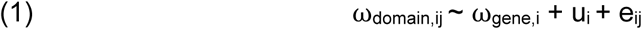

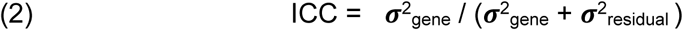

Here, ⍵_domain,ij_ is dN/dS of domain *j* in protein *i*, ⍵_gene,i_ is the gene-level dN/dS for protein *i*, u_i_ is the random effect, and e_ij_ is the residual error. 𝝈^2^ is the variance between genes, and 𝝈^2^ is the variance within genes (between domains).

To examine whether the relationship between domain and gene-level dN/dS is different among gene classes, we fit Equation 3 on a dataset containing all gene classes. In this model, the interaction between the gene class and ⍵_gene_ tests whether the slope between ⍵_gene_ and ⍵_domain_ is different across gene classes. We assessed the difference between ICC among classes using parametric bootstrapping with the bootMer function from the lme4 package.

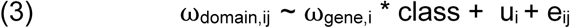

### Domain enrichment analyses

Protein domain enrichment was examined for pleiotropic, immune, and developmental genes relative to randomly selected (background) genes. For each dataset, each domain was counted once per protein. Enrichment analysis was performed using Fisher’s exact test on 2x2 contingency tables comparing the number of proteins with and without the domain in the target versus background sets, and P-values were adjusted using the Benjamini-Hochberg method. Odds ratios were calculated by adding a pseudo-count of 0.5 to avoid zero values (Equation 4):

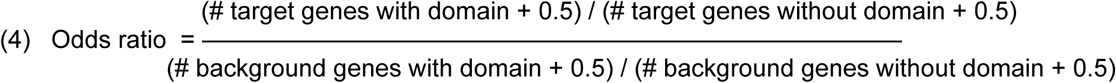

### Identifying conserved and radical replacements in genes and domains

To link dN/dS to possible structural change in the protein sequences, we calculated the ratio of radical to conserved (R/C) amino acid changes across the phylogenetic tree for both genes and domains. To this end, we extracted amino acid changes from the rst files generated by PAML across the phylogenetic tree for every gene and its domain(s). Each amino acid changes across a tree were classified either as radical or conservative based on the BLOSUM62 matrix (Henikoff and Henikoff 1992). Radical changes have negative scores in the matrix (less likely to occur), whereas conservative changes have positive scores (more likely to occur). To ensure consistency between R/C and dN/dS, we estimated R/C for unsaturated (dS < 3) genes and domains.

### Estimating ω_na_, ω_a_, and ⍺

For the four classes of genes (random, pleiotropic immune genes, non-pleiotropic immune genes, and developmental genes), we measured the non-adaptive substitution rate using MultiDFE (https://github.com/kousathanas/MultiDFE). This was done for three populations of *D. melanogaster* (Munich, Barcelona, and Raleigh) in the *Drosophila* Evolution over Space and Time (DEST) dataset (Kapun et al. 2021; Footprints of worldwide adaptation in structured populations of *Drosophila melanogaster* through the expanded DEST 2.0 genomic resource 2025). To this end, we first used degenotate (Mirchandani et al. 2024) to identify fourfold and zerofold degenerate sites and extracted their genomic coordinates for downstream analyses.

Next, we computed per-gene SFS from masked sync files at fourfold and zerofold degenerate sites, applying a minimum depth filter (sites were considered callable if A+T+C+G ≥ 10) and projecting allele frequencies to a fixed sample size of 20, similar to genome-wide SFS calculation in the DEST code repository. We next generated 100 bootstrap replicates of each of the three categories, summing the corresponding per-gene zerofold and fourfold SFS across genes, while rounding projected counts to preserve the total number of sites. The results were used as inputs to MultiDFE to measure ω_na_. For the gene-level analyses, we estimated the rate of adaptive substitution (ω_a_) and the proportion of adaptive substitution (⍺) using the dN/dS ratios (rate of zero to fourfold change) from the PopFly dataset in the iMKT R package (Murga-Moreno et al. 2019) following equations 10 and 11 in (Kousathanas and Keightley 2013).

For the domain-level SFS construction per gene, we blasted the CDS sequence of each domain to the *D. melanogaster* genome to identify its coordinates within the genome. Next, we removed redundancy caused by overlapping domains and also removed blast results that fall outside of the gene of interest (misidentified domains). Misidentification of the domain coordinates happened for five out of 1,267 random genes, four out of 3,047 developmental, one out of 454 non-pleiotropic immune genes, and none of the pleiotropic immune genes. For each gene, domain-level SFS is constructed by counting SNPs across the domain coordinates. For the domain-level analysis, we only estimated ω_na_ because dN/dS (the rate of zero to fourfold change) for the domain-level is not available.

### Gene expression across four species of *Drosophila*

To understand whether pleiotropy constraints the evolution of plasticity in gene expression, we downloaded the raw RNA-seq expression data from Salazar-Jaramillo et al. (2017) for control and infected larvae samples with parasitoid wasps for four species of *Drosophila* (*D.melanogaster*, *D. simulans, D. sechellia,* and *D. yakuba*). We chose samples for which RNA was extracted five hours post-exposure to parasites to capture the maximum immune response. Out of three lines of *D. melanogaster* (two wild-type and one evolved for resistance to wasps), we chose one wild-type line. Kallisto (Bray et al. 2016) was used to estimate gene expression by pseudo-alignment of RNA-seq reads to the following genome annotations: Release 6.54 for *D. melanogaster,* Release 103 for *D. simulans*, Release 102 for *D. yakuba,* and Release 101 for *D. sechellia*. For each gene, we summed counts across all transcripts. Differential gene expression analyses for four species were performed using DESeq2 (Love et al. 2014). We found log_2_ fold change values for orthologous genes across the four *Drosophila* species. For pleiotropic, immune genes, and random genes (as a background set), we used these log_2_ fold change values to perform principal component analysis (PCA). The Euclidean distance of each gene from the PCA centroid was used to define “central” genes, with thresholds ranging from the closest 5% to 50% of genes. The remaining genes were classified as “scattered”. We then compared the median dN/dS of central and scattered genes for unsaturated genes (dS < 3).

## Data Availability Statement

All scripts, data, and instructions for running the analyses are available here: https://github.com/danialasg74/Pleiotropic-Constraints-on-Immune-System-Evolution/tree/main.

## Author Contributions

A.T.T. and D.A. conceived and designed the study. D.A. performed the analyses and wrote the first draft. D.A. and A.T.T. contributed to the final draft.

## Acknowledgments

This work was supported by the National Institute of General Medical Sciences at the National Institutes of Health (grant number R35GM138007 to A.T.T.). AI was used under supervision to assist with writing some of the code.

