## Supplemental Material for "Pleiotropy constrains the evolution of immune system plasticity while promoting domain modularity"

### Supplementary material for the manuscript "*Pleiotropy constrains the evolution of immune system plasticity while promoting domain modularity*"

**Table S1. Parameter estimates from Equation 1, fitted separately for each gene class.**

Intercepts, fixed effects, and standard deviations for random effects are reported.

| **Gene Class** | **Intercept** | $\omega_{gene}$ | $Sd_{Protein}$ | $Sd_{residual}$ |
|---|---|---|---|---|
| Random | -0.03133 | 1.03372 | 0.06607 | 0.03264 |
| Immunity | -0.01339 | 0.89928 | 0.03603 | 0.04410 |
| Pleiotropy | -0.007772 | 0.626928 | 0.02101 | 0.04167 |
| Developmental | -0.006014 | 0.590275 | 0.02581 | 0.03276 |

**Table S2. Pairwise comparisons of estimated $\omega_{gene}$ effects.** Pairwise contrasts were obtained from the interaction term in Equation 3 using emtrends in R.

| **Contrasts** | **estimate** | **SE** | **Adjusted P-value** |
|---|---|---|---|
| Developmental - Immunity | -0.3102 | 0.0300 | <0.0001** |
| Developmental - Pleiotropic | -0.0309 | 0.0450 | 0.9025 |
| Developmental - Random | -0.4346 | 0.0236 | <0.0001** |
| Immunity - Pleiotropic | 0.2793 | 0.0490 | <0.0001** |
| Immunity - Random | -0.1244 | 0.0306 | 0.0003* |
| Pleiotropic - Random | -0.4037 | 0.0454 | <0.0001** |

**Table S3. Protein domain enrichment analysis in pleiotropic, immune, and developmental genes relative to a background set of randomly selected genes.** Odds ratios > 1 indicate enrichment, whereas odds ratios < 1 indicate depletion.

| Pleiotropic | Odds ratio | Immunity | Odds ratio | Developmental | Odds ratio |
|---|---|---|---|---|---|
| ADP-ribosylation factor family | 11.1 | C-type lysozyme/alpha-lactalbumin family | 20.6 | Helix-loop-helix DNA-binding domain | 10.5 |
| Protein kinase domain | 10.1 | Gamma-thionin family | 39.9 | Fibronectin type III domain | 24.9 |
| Ras family | 7.87 | CD36 family | 45.4 | Homeodomain | 63.0 |
| Protein tyrosine and serine/threonine kinase | 10.1 | N-acetylmuramoyl-L-alanine amidase | 50.8 | Protein kinase domain | 4.54 |
| Ras of Complex, Roc, domain of DAPkinase | 10.1 | Stress-inducible humoral factor Turandot | 45.4 | Insect cuticle protein | 27.4 |
| | | | | Protein tyrosine and serine/threonine kinase | 4.54 |
| | | | | Trypsin | 0.202 (depletion) |
| | | | | Protein of unknown function (DUF1091) | 0.0267 (depletion) |
| | | | | 7tm Chemosensory receptor | 0.0411 (depletion) |

**Table S4. ICC values for domains within proteins containing at least one kinase domain across four gene classes.** Hare, n represents the number of genes in each class.

| Group | ICC |
|---|---|
| Pleiotropic (n = 40) | 0.05941474 |
| Immunity (n = 9) | 0.596376 |
| Developmental (n = 155) | 0.7758814 |
| Random (n = 25) | 0.4032117 |

**Table S5. Medians and adjusted p-values for comparisons are presented in Fig.4 panels A-D.**

| **Genes dN/dS** | | | | | |
|---|---|---|---|---|---|
| **Class** | **Median** | **Recognition** | **Signaling** | **Effector** | **Anti-viral** |
| **Recognition** | 0.094 | - | - | - | - |
| **Signaling** | 0.060 | 0.0051* | - | - | - |
| **Effector** | 0.089 | 0.7055 | 0.0221* | - | - |
| **Anti-viral** | 0.114 | 0.9320 | 0.1480 | 0.9320 | - |
| **Other** | 0.042 | 0.0072* | 0.4866 | 0.0182* | 0.1480 |
| **Domains dN/dS** | | | | | |
| **Class** | **Median** | **Recognition** | **Signaling** | **Effector** | **Anti-viral** |
| **Recognition** | 0.036 | - | - | - | - |
| **Signaling** | 0.014 | 3.4e-07*** | - | - | - |
| **Effector** | 0.037 | 0.175 | 2.2e-10*** | - | - |
| **Anti-viral** | 0.077 | 0.044* | 2.6e-06*** | 0.166 | - |
| **Other** | 0.025 | 0.428 | 2.5e-06*** | 0.015* | 0.015* |
| **Genes R/C** | | | | | |
| **Class** | **Median** | **Recognition** | **Signaling** | **Effector** | **Anti-viral** |
| **Recognition** | 0.600 | - | - | - | - |
| **Signaling** | 0.713 | 0.430 | - | - | - |
| **Effector** | 0.546 | 0.255 | 0.044* | - | - |

| | | | | | |
|---|---|---|---|---|---|
| **Anti-viral** | 0.547 | 0.255 | 0.117 | 0.846 | - |
| **Other** | 0.594 | 0.341 | 0.063 | 0.739 | 0.640 |
| **Domains R/C** | | | | | |
| **Class** | **Median** | **Recognition** | **Signaling** | **Effector** | **Anti-viral** |
| **Recognition** | 0.754 | - | - | - | - |
| **Signaling** | 0.724 | 0.78206 | - | - | - |
| **Effector** | 0.538 | 2.8e-05*** | 2.2e-06*** | - | - |
| **Anti-viral** | 0.551 | 0.00267** | 0.00092*** | 0.59127 | - |
| **Other** | 0.554 | 7.8e-07*** | 4.2e-08*** | 0.72219 | 0.94289 |

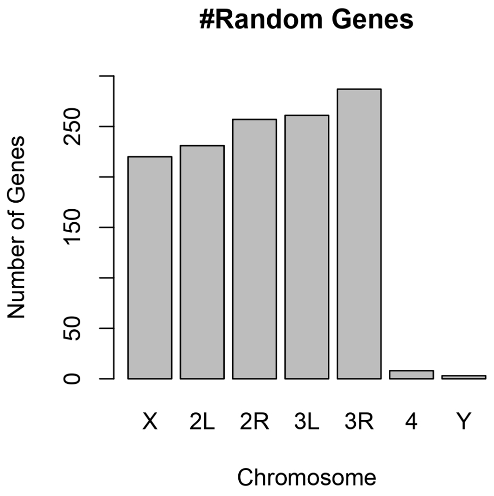


**FigS1. Number of randomly selected genes across the *Drosophila* genome.** The X-axis shows the chromosomes, and the Y-axis shows the number of randomly selected genes per chromosome.

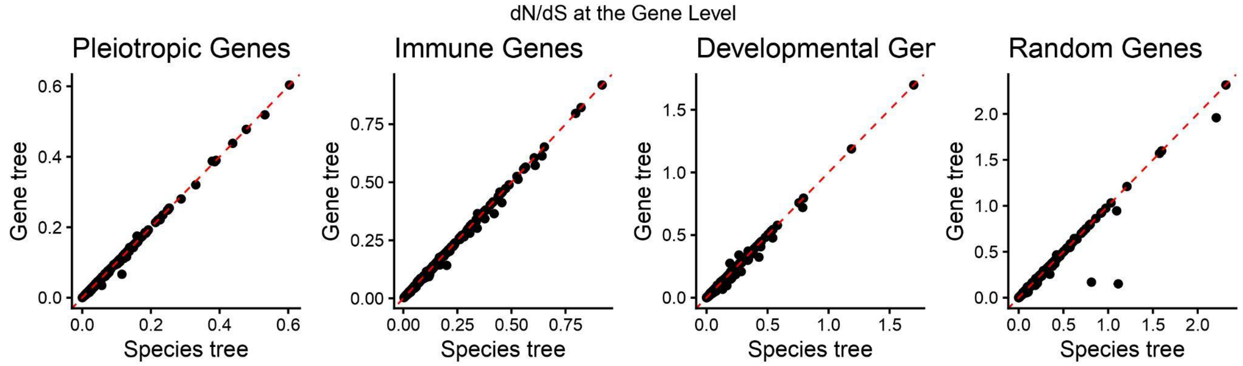


**FigS2. dN/dS estimates are highly consistent between the species tree (X-axis) and the gene tree (Y-axis).** The red dashed lines show the y = x line.

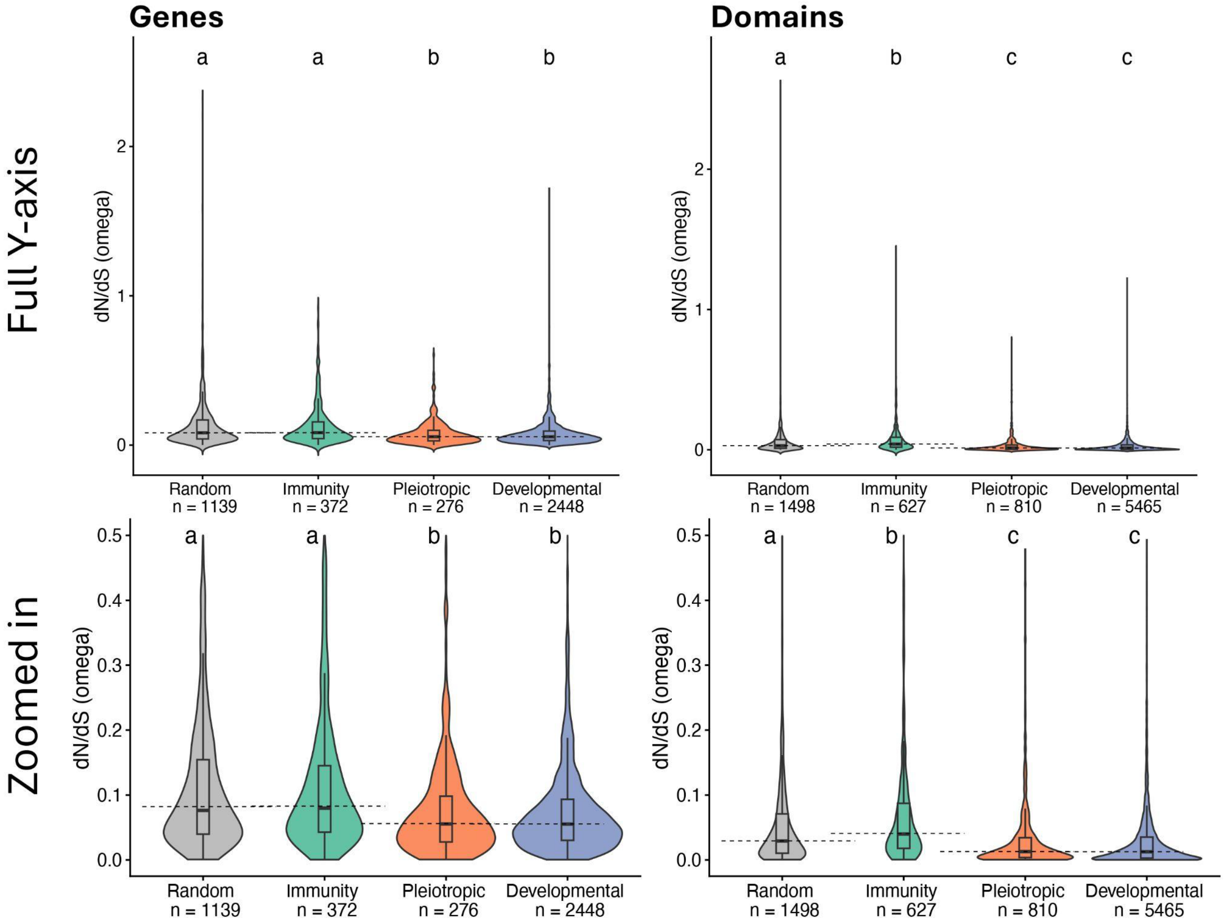


**FigS3. Non-pleiotropic immune genes evolve faster than pleiotropic immune genes and developmental genes.** The dN/dS values are plotted for genes (left) and domains across three groups of genes (immunity = non-pleiotropic, pleiotropic = pleiotropic immune genes, and

developmental). Groups sharing a letter are not significantly different (Wilcoxon test). The bottom panel shows the zoomed-in version of the top panel. To improve visualization, medians are extended with black dashed lines to facilitate comparison.

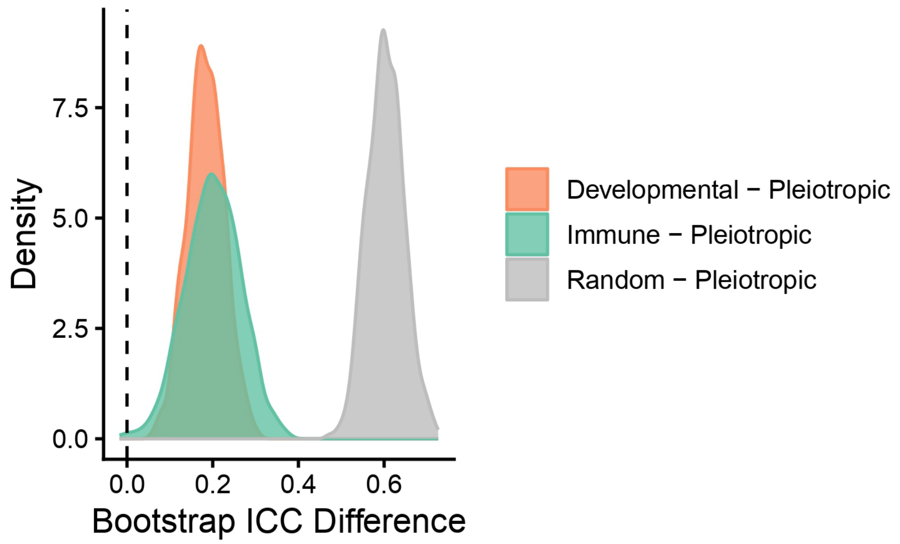


**FigS4. Parametric bootstrapping (n=1000) using Equation 1 shows Pleiotropic genes have smaller ICC than other gene classes in nearly 100% of comparisons.** The X-axis shows the difference in bootstrap values, and the Y-axis is the density. The zero difference is shown with a vertical dashed line. Positive values on the X-axis indicate a smaller ICC for the pleiotropic group of genes.

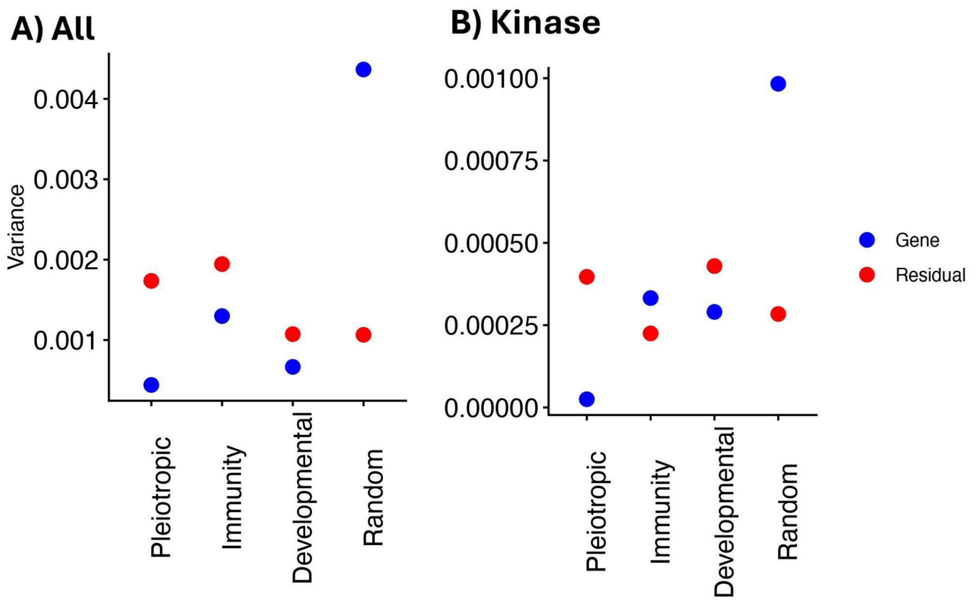


**FigS5. Low intra-class correlation coefficient (ICC) for dN/dS estimates in pleiotropic genes is associated with low between-gene variance and elevated within-gene (between domains) variance.** The variance components, namely, the gene and residual variance, are shown. The left panel shows the analysis for all genes, while the right panel shows it for domains with at least one kinase domain.

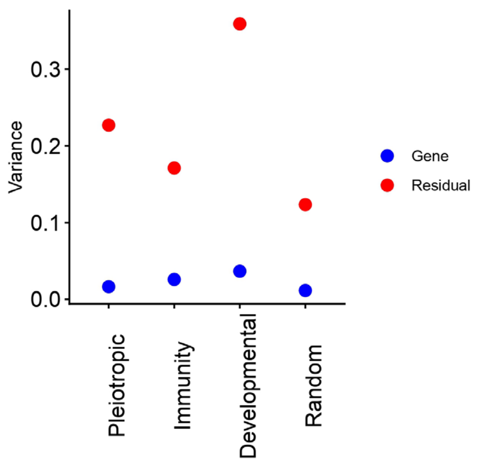

**FigS6. Low intra-class correlation coefficient (ICC) for R/C estimates across gene classes is associated with low between-gene variance.** The variance components, namely, the gene and residual variance, are shown.

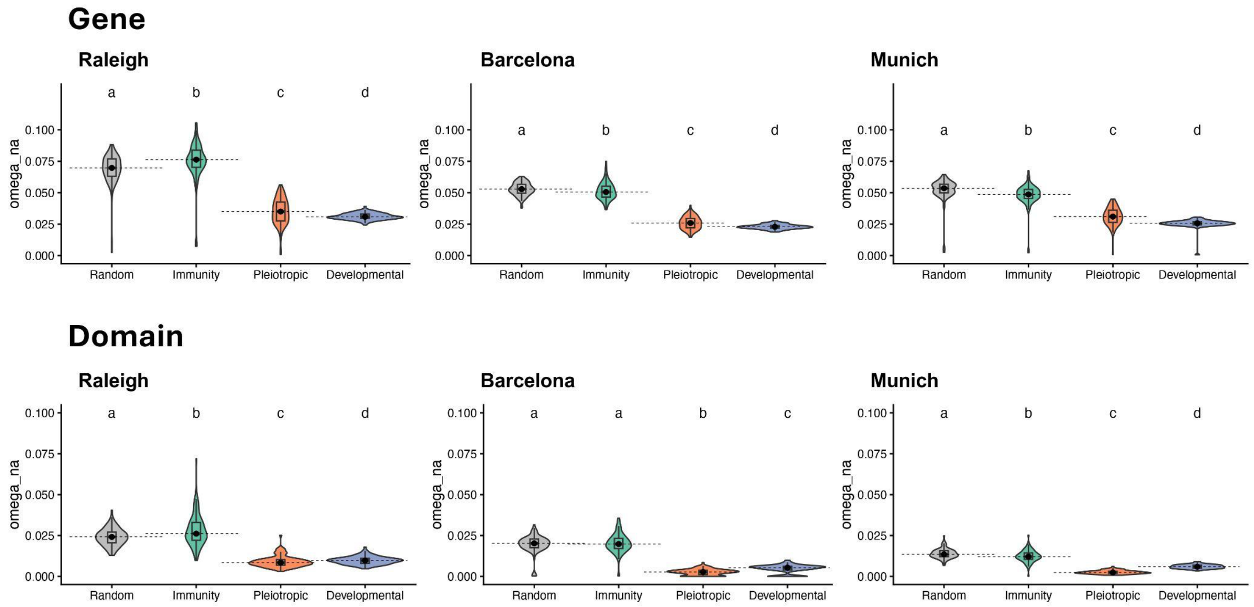


**FigS7. Pleiotropic immune genes have the second-lowest rate of non-adaptive substitutions at the gene level and the lowest at the domain level.** The Y-axis shows the rate of non-adaptive substitution ($\omega_{na}$) across three classes of genes (X-axis) for the Raleigh population. The top row shows the results at the gene level and the bottom row at the domain level. Each column shows $\omega_{na}$ values for a population of *D. melanogaster.* Groups sharing a letter are not significantly different (Wilcoxon test). To improve visualization, medians are extended with black dashed lines to facilitate comparison.

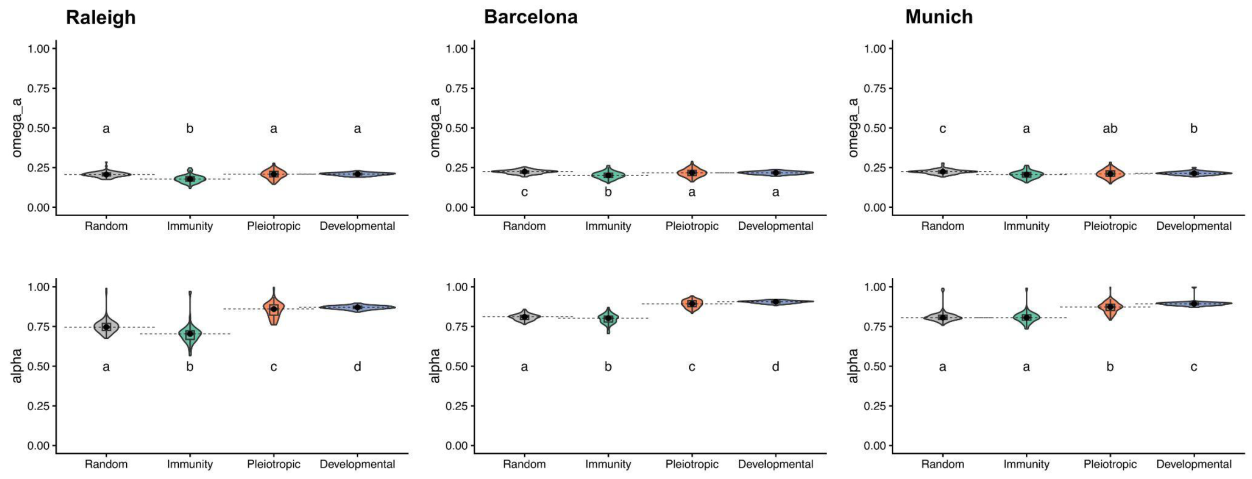


**FigS8. Across three populations of D. melanogaster (columns), pleiotropic immune genes have a higher rate and proportion of adaptive substitutions compared to non-pleiotropic immune genes, and are closer to developmental genes.** The top row shows the rate of adaptive substitutions ($\omega_a$), while the bottom row shows the proportion of adaptive substitutions ($\alpha$). Groups sharing a letter are not significantly different (Wilcoxon test). To improve visualization, medians are extended with black dashed lines to facilitate comparison.

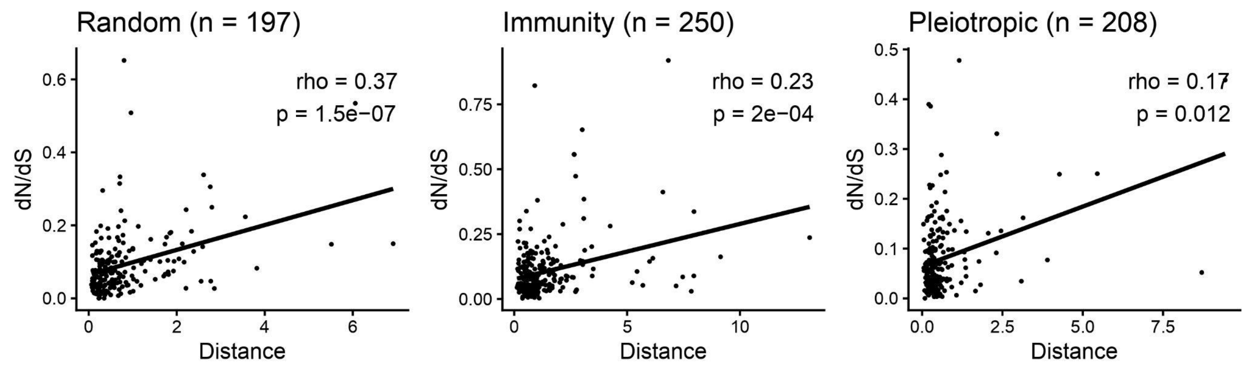


**FigS9. Weak but significant positive correlation between variability in log2 fold changes following exposure to parasitoid wasps and dN/dS for orthologous genes across four species of *Drosophila.*** The strength of correlation ($\rho$) is strongest for random genes, moderate

for non-pleiotropic immune genes, and lowest in pleiotropic ones. The dN/dS values here are only reported for dS<3.

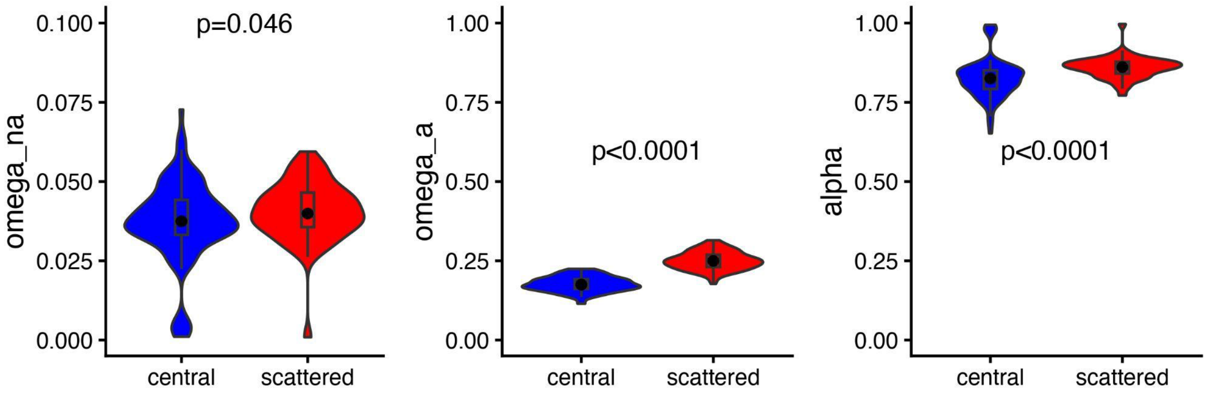


**FigS10. The higher dN/dS in scattered compared to central non-pleiotropic immune genes at 50% threshold is associated with adaptive evolution.** The left panel shows the rate of non-adaptive mutations, which is slightly higher in scattered genes. The central and right panels show the rate of adaptive substitutions and the proportion of adaptive substitutions, which are significantly greater in scattered genes.
